# rrQNet: protein contact map quality estimation by deep evolutionary reconciliation

**DOI:** 10.1101/2022.01.20.477079

**Authors:** Rahmatullah Roche, Sutanu Bhattacharya, Md Hossain Shuvo, Debswapna Bhattacharya

## Abstract

Protein contact maps have proven to be a valuable tool in the deep learning revolution of protein structure prediction, ushering in the recent breakthrough by AlphaFold2. However, self-assessment of the quality of predicted structures are typically performed at the granularity of 3D coordinates as opposed to directly exploiting the rotation- and translation-invariant 2D contact maps. Here we present rrQNet, a deep learning method for self-assessment in 2D by contact map quality estimation. Our approach is based on the intuition that for a contact map to be of high quality, the residue pairs predicted to be in contact should be mutually consistent with the evolutionary context of the protein. The deep neural network architecture of rrQNet implements this intuition by cascading two deep modules—one encoding the evolutionary context and the other performing evolutionary reconciliation. The penultimate stage of rrQNet estimates the quality scores at the interacting residue-pair level, which are then aggregated for estimating the quality of a contact map. This design choice offers versatility at varied resolutions from individual residue pairs to full-fledged contact maps. Trained on multiple complementary sources of contact predictors, rrQNet facilitates generalizability across various contact maps. By rigorously testing using publicly available datasets and comparing against several in-house baseline approaches, we show that rrQNet accurately reproduces the true quality score of a predicted contact map and successfully distinguishes between accurate and inaccurate contact maps predicted by a wide variety of contact predictors. The open-source rrQNet software package is freely available at https://github.com/Bhattacharya-Lab/rrQNet.

## INTRODUCTION

Deep neural networks have led to a paradigm shift in protein structure prediction^1–5^. Most notably, the AlphaFold2 method from DeepMind^5^ has demonstrated unprecedented performance level in the 14th edition of the Critical Assessment of Structure Prediction (CASP14) experiment^6^ through successful application of deep neural networks, paving the way towards highly accurate prediction of protein structural models at proteome-wide scale^7^. The core of AlphaFold2 consists of a neural embedding of the evolutionary profile coupled with pairwise relations of the various amino acid residues in the protein. This embedding is used to predict the inter-residue interactions between the amino acids by a collection of deep neural network modules trained end-to-end, leading to the final predicted three-dimensional (3D) structure. Inter-residue interactions can be represented as a binary two-dimensional (2D) matrix, also known as a contact map, which distinguishes interacting residue pairs from the non-interacting ones using a predefined distance threshold such as 8Å. Typically, if the spatial positioning of the C_β_ (C_α_ in case of Glycine) atom of a residue i lies within 8Å of the C_β_ (C_α_ in case of Glycine) atom of residue j, the residue pair (i,j) is considered to be an interacting pair (contact). A contact map has the advantage of being invariant to rotations and translations, while providing a 2D representation of a protein 3D structure. Intuitively, the choice of the distance threshold affects the granularity of a contact map, and the use of multiple distance thresholds results in finer-grained binned distance distributions. Regardless of their resolutions, contact maps encode inter-residue spatial proximity information that can be turned into Cartesian coordinates, leading to the predicted structure^8^. As such, a 2D contact map can be considered as a transitional state in the mapping from the 1D amino acid sequence to the 3D structure of proteins.

In order for protein structure prediction to be practically useful, a reliable self-assessment of the prediction is critically important^9,10^. Such a calibrated self-assessment of the quality of the prediction is an integral part of the AlphaFold2 system, which provides intrinsic model accuracy estimates in the form of predicted local-distance difference test (pLDDT) and predicted global superposition metric template modeling score (pTM). However, self-assessment of the quality of structure prediction using the state-of-the-art protein structure prediction systems typically rely on 3D coordinates as opposed to exploiting the residue pair representation captured by a 2D contact map. That is, the granularity of protein quality estimation is typically a predicted 3D structure rather than a 2D contact map.

Performing self-assessment at the granularity of 2D contact maps instead of 3D structures has several advantages. First, being invariant to rotations and translations, contact maps and their estimated confidence scores can be seamlessly integrated into geometric attention-based deep learning models while implicitly ensuring invariance^5^. Second, the estimated quality score of a 2D contact map can be integrated into the loss function of an end-to-end structure prediction system. Third, such an integration enables error feedback^11^ to propagate through the deep network for iterative optimization of the contact maps, similar to recycling updates performed in AlphaFold2. Finally, a reliable quality estimate of predicted contact maps can inform contact-driven homology detection and reconstruction methods^8,12–19^.

How can we get the representational benefits of self-assessment of the prediction at the 2D contact map level, without necessarily turning the contact map into 3D coordinates? Here we explore a potential solution of estimating the contact map quality by *deep evolutionary reconciliation*. Our approach is based on the intuition that for a contact map to be of high quality, the residue pairs predicted to be in contact should be mutually consistent with the evolutionary context encoded in the multiple sequence alignment of the target protein sequence. The deep neural network architecture of rrQNet directly implements this intuition by cascading two deep modules—one encoding the evolutionary context and the other performing evolutionary reconciliation. The penultimate stage in rrQNet is a rough estimate of the predicted confidence score at the level of interacting residue pairs, which is subsequently aggregated for scoring a predicted contact map. We rigorously test rrQNet on publicly available datasets by comparing against several baseline approaches and conducting pseudo-blind assessment using protein targets from the latest rounds of CASP experiments for several hundred predicted contact maps generated by a wide variety of contact predictors. The experimental results show that our method can accurately reproduce the true quality score of a predicted contact map and can successfully distinguish between accurate and inaccurate contact prediction. Our generalizable deep learning model is versatile for confidence estimation at varying granularities from individual residue-pair to full-fledged contact map. The open-source rrQNet software package is freely available at https://github.com/Bhattacharya-Lab/rrQNet.

## MATERIALS AND METHODS

### Model Architecture

As shown in **Figure 1**, our deep learning model for protein contact map quality estimation consists of two major modules, each being a residual neural network (ResNet)^20^. The first module is the *evolutionary module*, which conducts a series of 2D convolutional transformations on the evolutionary context extracted from the covariational signal encoded in the multiple sequence alignment (MSA). The output of the evolutionary module is then converted to a 2D matrix and then fed into the second module together with the predicted contact maps. The second module is the *reconciliation module*, which performs a series of 2D convolutional transformations of the evolutionary context-integrated contact maps. Finally, the output of the reconciliation module is utilized to predict the correctness of contacts at the residue pair level, which can be subsequently aggregated for estimating the overall quality of the predicted contact map.

**Figure 1.**
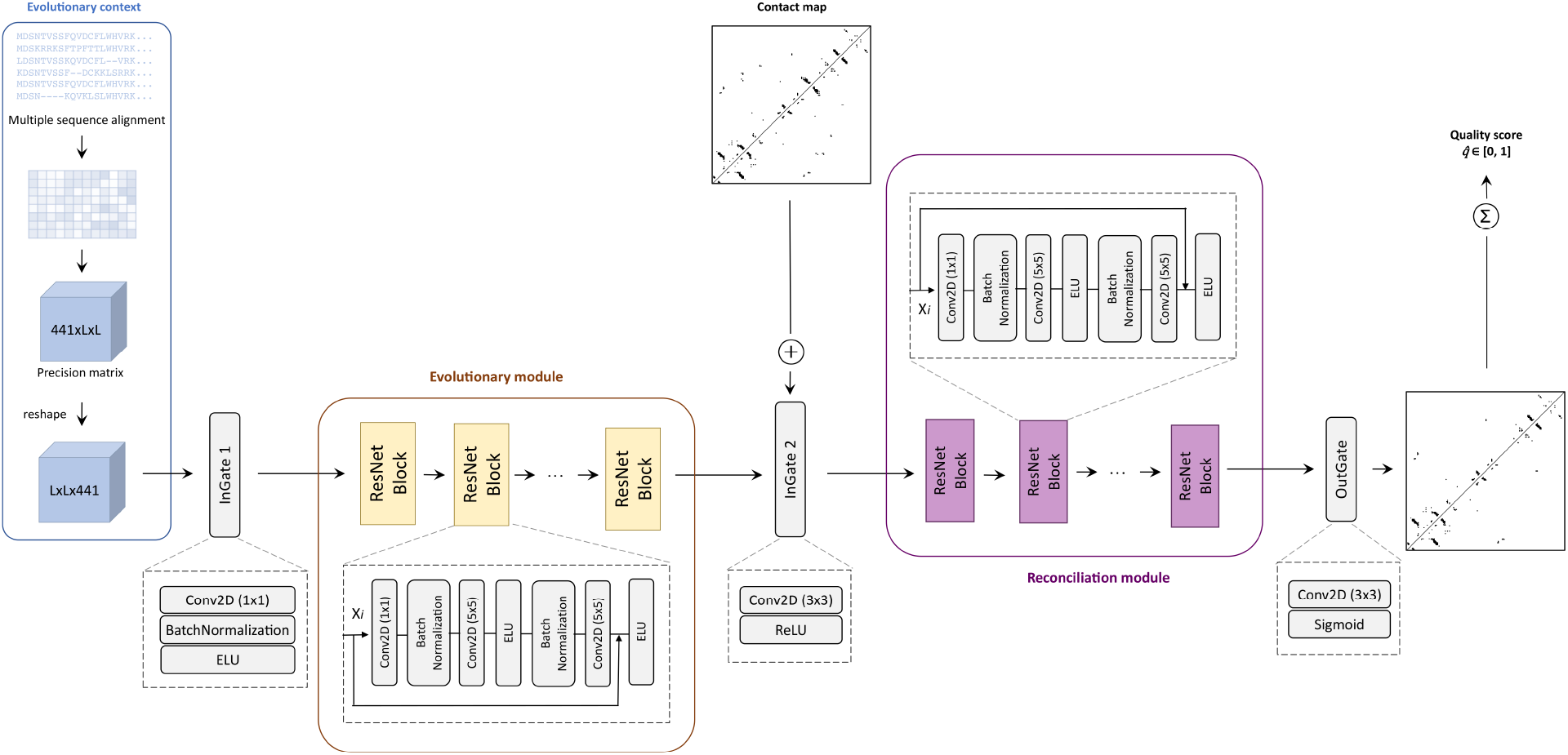
rrQNet architecture overview. Left to right: The evolutionary context extracted from the covariational signal encoded in the multiple sequence alignment (MSA) is fed to the evolutionary module, which conducts a series of 2D convolutional transformations. The output of the evolutionary module is then converted to a 2D matrix and then fed into the reconciliation module together with the predicted contact map to perform evolutionary reconciliation. Finally, the output of the reconciliation module is utilized to predict the correctness of contacts at the residue pair level, which can be subsequently aggregated for estimating the overall quality of the full-fledged contact map.

#### Inputs for training and inference

The inputs to rrQNet comprise of a predicted contact map and multiple sequence alignment (MSA) for the target protein sequence. MSA is generated by searching against diverse sequence sources after merging sequences from whole-genome sequence databases and from metagenome database^21^. From the MSA, an inverse co-variation matrix (or precision matrix)^22^ is generated which captures dependent conditional correlations among pairwise variables. A precision matrix contains L×L blocks (L is the length of the protein sequence) and each block contains 21×21 matrix for direct couple correlation for a particular pair of residues^23^ (i.e., having dimension of 21×21=441 for each pair of residues). We reshape the 441×L×L precision matrix to L×L×441 before feeding into the evolutionary module. During training, the input MSA is used for generating a diverse array of contact maps using seven different methods: CCMpred^24^, FreeContact^25^, PSICOV^26^, MetaPSICOV^27^, DeepConPred2^28^, DNCON2^29^, and ResPRE^23^. CCMpred is a pseudo-likelihood maximization (PLM)-based approach^30,31^ for contact prediction optimized for performance. FreeContact is a speed-optimized implementation of EVfold-mfDCA^12^. PSICOV uses sparse inverse co-variation estimation^32^ for predicting protein contact map. MetaPSICOV is a two-stage contact predictor that employs a combination of approaches for obtaining MSA-derived co-variation signal and various statistics computed from the MSA for contact prediction. DeepConPred2 is an improved re-optimized version of the deep learning-based approach DeepConPred^33^. DNCON2 employs a two-level deep convolutional neural network for contact prediction. ResPRE leverages residual neural network architecture coupled with precision matrix to predict contact maps. All methods are locally installed and run with default parameter settings to generate contact maps and top L contacts (L is the length of protein sequence) are fed to the reconciliation module. Following CASP standard of contact map assessment and to ensure a fair comparison between the contact predictors, we consider top L contacts for all methods. Our approach can be easily adapted for any number of contact pairs including all residue pairs. During inference, rrQNet takes only one contact map as input predicted by any method (including but not limited to the seven contact predictors used for training) along with the multiple sequence alignment.

#### Evolutionary module

The evolutionary module is preceded by a gate (InGate 1) comprising of a 2D convolutional layer with a kernel size of 1×1, batch normalization, and a nonlinear transformation called exponential linear unit (ELU). The input to this gate is the reshaped precision matrix of size L×L×441, which is transformed to a size of L×L×64, using a 1×1 convolutional layer with a filter size of 64. This input gate is followed by a stack of 55 residual neural network (ResNet) blocks. Each residual block in this ResNet stack consists of three 2D convolutional layers with a filter size of 64. The number of ResNet blocks is determined through validation on the independent CASP11 dataset to avoid overfitting, as discussed later. The first convolutional layer with kernel size 1×1 is used to restore the dimensionality of the feature vector and is followed by batch normalization. The second convolutional layer with kernel size of 5×5 is followed by ELU activation unit and batch normalization. The third convolutional layer with kernel size of 5×5 is the output layer of a ResNet block having a shortcut connection to the input layer for skipping the intermediate layers. This output propagates to the next ResNet block through a nonlinear transformation via the ELU unit.

#### Reconciliation module

Similar to the evolutionary module, the reconciliation module is preceded by a gate (InGate 2) consisting of a 2D convolutional layer having a kernel size of 3×3 and filter of 1, followed by a different transformation called rectified linear unit (ReLU) to transform the size to L×L×1, before concatenating with the input contact map. The reconciliation module leverages a stack of 40 residual neural network blocks having architecture similar to the evolutionary module determined through validation on the independent CASP11 dataset to avoid overfitting, as discussed later.

#### Quality estimation

The reconciliation module is followed by a gate (OutGate) comprising of a 2D convolutional layer having a kernel size of 3×3 followed by sigmoidal transformation, whose output leads to a 2D matrix of predicted confidence score at the level of a pair of interacting amino-acid residues from the input contact map. The 2D matrix of predicted confidence score is then subsequently aggregated for estimating the final quality score 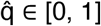 of the full-fledged input contact map as: 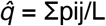, where L is the number of residues, pij is 1 if the average predicted likelihood of residue pair (i,j) and the residue pair (j,i) is greater than the threshold of 0.5, and 0 otherwise. As such, 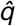 is analogous to the precision of the input contact map.

### Training and implementation details

Our method is implemented using Keras with TensorFlow^34^ backend and trained using Quadro RTX 5000 GPU with 16GB memory. We train our model for 50 epochs and use checkpoint for saving the best model on each epoch. Models are trained on a batch size of 2, shuffling the training data, using ADAM optimizer^35^ with a learning rate of 0.001 and binary cross-entropy loss function. While training, we crop (or pad) the input dimension to 256, but the trained model can estimate the quality score for any input length.

### Datasets for training and performance evaluation

For training the deep learning model, we use a total of 3,073 contact maps of varying qualities predicted by seven different contact prediction methods previously discussed using a non-redundant dataset of 439 proteins including 421 protein targets from the DeepCov^36^ dataset and 18 free modeling (FM) target domains from the 7th through 10th editions of the Critical Assessment of Protein Structure Prediction (CASP7-CASP10) experiments. We consider protein chains having no more than 35% pairwise sequence identity to remove redundancy in the training set.

For performance evaluation, we use a test set of 280 contact maps predicted by the seven locally installed contact prediction methods for 40 FM target domains with publicly available experimental structures from the 12th and 13th editions of CASP experiments (CASP12 and CASP13) to perform benchmark assessment; and 660 contact maps predicted by 30 fully automated server predictors participating in the contact prediction category of the 14th edition of CASP (CASP14) for 22 FM targets to perform pseudo-blind assessment. Additionally, 210 contact maps predicted by the locally installed seven contact prediction methods for 30 FM target domains from the 11th edition of CASP (CASP11) are used as a validation set. Both the test and validation sets are structurally independent from the training set, having TM-align score < 0.5^37^. For the benchmark assessment on CASP12 and CASP13 sets as well as validation on CASP11 set, we run the seven locally installed contact predictors with default parameter settings using multiple sequence alignments (MSA) generated by searching against diverse sequence sources including whole-genome sequence databases (Uniclust30^38^, UniRef90^39^) and metagenome database (Metaclust^40^) using DeepMSA^21^ pipeline with default parameters. Top L predicted contacts (L is the length of the protein sequence) are then fed into rrQNet for contact map quality estimation. For CASP14 pseudo-blind assessment, we download the contact predictions submitted by the server predictors directly from the CASP website and feed them into rrQNet along with the corresponding MSAs for quality estimation.

### Comparison against other baseline approaches

To evaluate the effectiveness of our method in the absence of other competing approaches, we compare its performance against three in-house baseline contact quality estimation methods: consensus, shallow neural network, and naïve Bayes. The consensus-based baseline estimates the quality of a predicted interacting residue-pair by a contact predictor based on the average likelihood values of the predictions made for that residue-pair by the other contact predictors. That is, the average estimated confidence score across the other contact predictors for a residue-pair is its consensus-based quality score. Following this approach, we calculate a consensus score of each predicted contact in a contact map to estimate the quality at the residue-pair level and subsequently aggregate the resulting 2D matrix containing the consensus residue-pair confidence scores for estimating the final quality score of the full-fledged input contact map. The shallow neural network baseline consists of two fully connected layers. The precision matrix and the input contact map are concatenated and passed to the first layer having ten neurons. The second layer consists of one neuron followed by a sigmoid activation function whose output leads to a 2D matrix of predicted confidence score at the residue-pair level for subsequent aggregation and the final quality estimation of the full-fledged input contact map. The naïve Bayes baseline implements a Gaussian naïve Bayes using the precision matrix concatenated with the input contact map for contact map correctness estimation of at the residue-pair level to be subsequently aggregated for the final quality estimation of the full-fledged input contact map. For a fair comparison against the controls, the shallow neural network and naïve Bayes baselines are trained on the same training dataset as used in rrQNet.

### Evaluation metrices

We employ twofold evaluation criteria for the performance assessment: (i) capability to reproduce the true quality score of a predicted contact map and (ii) capability to distinguish between accurate and inaccurate contact prediction. Ground truth is quantified by computing the precision of a predicted contact map against the contact map derived directly from the experimental 3D structure as: precision = TP/ (TP+FP), where TP (true positive) is the number of correctly predicted contacts and FP (false positive) is the number of incorrectly predicted contacts. For the first evaluation criterion, we calculate Pearson (r), Spearman (ρ), and Kendal’s tau (τ) correlation coefficients between the estimated contact map quality and ground truth precision. We compute the per-target average correlation coefficients (Per-target average r, ρ, τ) by pooling together all predicted contact maps for a specific protein target, as well as the overall global correlation coefficients (Overall global r, ρ, τ) by considering all predicted contact maps for all targets. The higher the correlation is, the better the performance is. The second evaluation criterion is measured by the area under the receiver operating characteristic curve (ROC) with a ground truth precision cutoff of 0.6. The AUC (Area Under Curve) assesses how well the predicted quality score may distinguish accurate prediction from inaccurate ones. The larger the area under the curve (AUC) of ROC, the better the capability is to distinguish between accurate and inaccurate contact prediction. We also consider per-node strength correlation between the predicted and native contact maps, which is a graph-based evaluation metric introduced in a recent CASP14 contact and distance prediction assessment^41^. A contact map can be considered as a graph, in which the nodes represent the residues and the edges represent the pair-wise contacts between the residues. In the context of a contact map, the per-node strength of a node i can be determined simply by summing up its adjacent edges. We use the correlation between the per-node strength of a predicted contact map and the per-node strength of the native contact map to evaluate the predicted contact map, thus applying a topological graph-based measure.

## RESULTS

### Quality estimation performance on CASP12 and CASP13 benchmark sets

**Table 1** reports the performance of rrQNet as well as the baseline approaches in reproducing the true quality scores of predicted contact maps for 154 predicted contact maps for 22 FM target domains from CASP12 and 126 predicted contact maps for 18 FM target domains from CASP13. rrQNet consistently attains high positive correlations between the estimated contact map quality and ground truth precision that are significantly better than all the baseline approaches. For instance, rrQNet attains the highest per-target average Pearson correlation (r) > 0.85, Spearman correlation (ρ) > 0.8, and Kendal’s tau correlation (τ) > 0.65; as well as the highest global Pearson and Spearman correlations ∼0.8 (*p*-value < 3.9e-28) and Kendal’s tau correlation ∼0.6 (*p*-value < 8.85e-23).

**Table 1.** rrQNet performance in reproducing true quality scores of predicted contact maps on CASP12 and CASP13 targets compared to our in-house baseline approaches implementing consensus, shallow neural network (NN), and naïve Bayes (NB).

| Dataset | Method | Avg. $r$ | Avg. $\rho$ | Avg. $\tau$ | Global $r$ | Global $\rho$ | Global $\tau$ |
| --- | --- | --- | --- | --- | --- | --- | --- |
| CASP 12 | rrQNet | **0.87** | **0.80** | **0.65** | **0.81** | **0.79** | **0.60** |
|  | Consensus | -0.49 | -0.51 | -0.41 | 0.72 | 0.72 | 0.53 |
|  | NN | 0.09 | 0.09 | 0.11 | 0.75 | 0.74 | 0.53 |
|  | NB | 0.75 | 0.69 | 0.51 | -0.11 | -0.06 | -0.04 |
| CASP 13 | rrQNet | **0.90** | **0.84** | **0.71** | **0.81** | **0.79** | **0.59** |
|  | Consensus | -0.54 | -0.46 | -0.39 | 0.49 | 0.55 | 0.40 |
|  | NN | 0.14 | 0.12 | 0.14 | 0.58 | 0.57 | 0.39 |
|  | NB | 0.64 | 0.57 | 0.44 | -0.23 | -0.22 | -0.14 |

*Note:* Values in bold represent the best performance.

Consistently better performance of rrQNet compared to the baselines demonstrates the effectiveness of the deep neural network architecture adopted in rrQNet. For the baseline approaches, it is interesting to note the performance tradeoff between the per-target average correlations and the global correlations. For example, the consensus-based baseline performs reasonably well in terms of global correlations, but attains negative per-target average correlations for both CASP12 and CASP13. The shallow neural network baseline achieves weak positive per-target average correlations on both CASP12 and CASP13, but shows reasonable performance in terms of global correlations. On further inspection, we find that the consensus and the shallow neural network baselines often misclassify most of the true positive contacts and estimate low-quality scores for the full-fledged contact maps when the ground truth quality scores are in fact much higher, leading to poor per-target average correlations. For example, the consensus approach misclassifies most of the true positive contacts for contact map predicted by ResPRE for the target T0864-D1, resulting in a final quality score of 0.04 (lowest in the pool of estimated quality scores for target T0864-D1), whereas the ground truth score is 0.9 (highest in the pool of ground truth) scores for the target T0864-D1). On the other hand, for the contact map predicted by PSICOV for target T0864-D1, the final quality estimation score by consensus approach is 0.1 (highest in the pool of estimated quality scores for the target T0864-D1), whereas the ground truth score is 0.28 (second lowest in the pool of ground truth scores for the target T0864-D1). As such, the consensus-based baseline results in negative correlations (r = -0.92, ρ = -0.88, τ = -0.78) for the target T0864-D1. Similarly, for the target T0968s1-D1, the consensus-based method misses most of the true positive contacts predicted by ResPRE and estimates ResPRE as lower quality (consensus-based quality estimation score is 0.27, whereas the ground truth is 0.82) than FreeContact (consensus-based quality estimation score is 0.29, whereas the ground truth is 0.35), leading to negative correlations (r = -0.91, ρ = -0.82, τ = -0.69). Likewise, the shallow neural network baseline misclassifies most of the true positive contacts for contact map predicted by MetaPSICOV for the target T0862-D1, resulting in negative correlations (r = -0.83, ρ = - 0.68, τ = -0.45). By contrast, the naïve Bayes baseline attains reasonable per-target average correlations, but negative global correlations. Closer inspection reveals that the negative global correlations are caused by the naïve Bayes baseline frequently estimating higher quality scores (>0.7) for contact maps predicted by ResPRE for several targets including T0859-D1, T0862-D1, T0863-D1, T0990-D2, and T0990-D3 when the ground truth scores are low (<0.3); and lower quality scores (<0.4) for targets including T0864-D1, T0912-D3, T0968s2-D1, and T1022s1-D1 when the ground truth scores are high (>0.8). While the baseline methods exhibit performance tradeoff, rrQNet consistently delivers high performance on both global and per-target correlations. That is, rrQNet is able to accurately reproduce the true contact map quality score across all assessment metrics.

To investigate the capability of rrQNet against the baseline approaches in distinguishing between accurate and inaccurate contact prediction, we perform ROC analysis using all 280 contact predictions from combined CASP12 and CASP13 set containing 40 FM target domains. **Figure 2** shows the rrQNet ROC curve with a high AUC value of 0.94, which is better than all the baseline methods including consensus, shallow neural network and naïve Bayes. Except for naïve Bayes, which has very low AUC value since it frequently estimates higher quality scores for when the ground truth scores are low and lower quality scores when the ground truth scores are high, the other baselines achieve reasonable AUC values, albeit much lower than rrQNet. Overall, the results demonstrate that rrQNet attains superior performance in separating accurate and inaccurate contact prediction compared to the baseline approaches.

**Figure 2.**
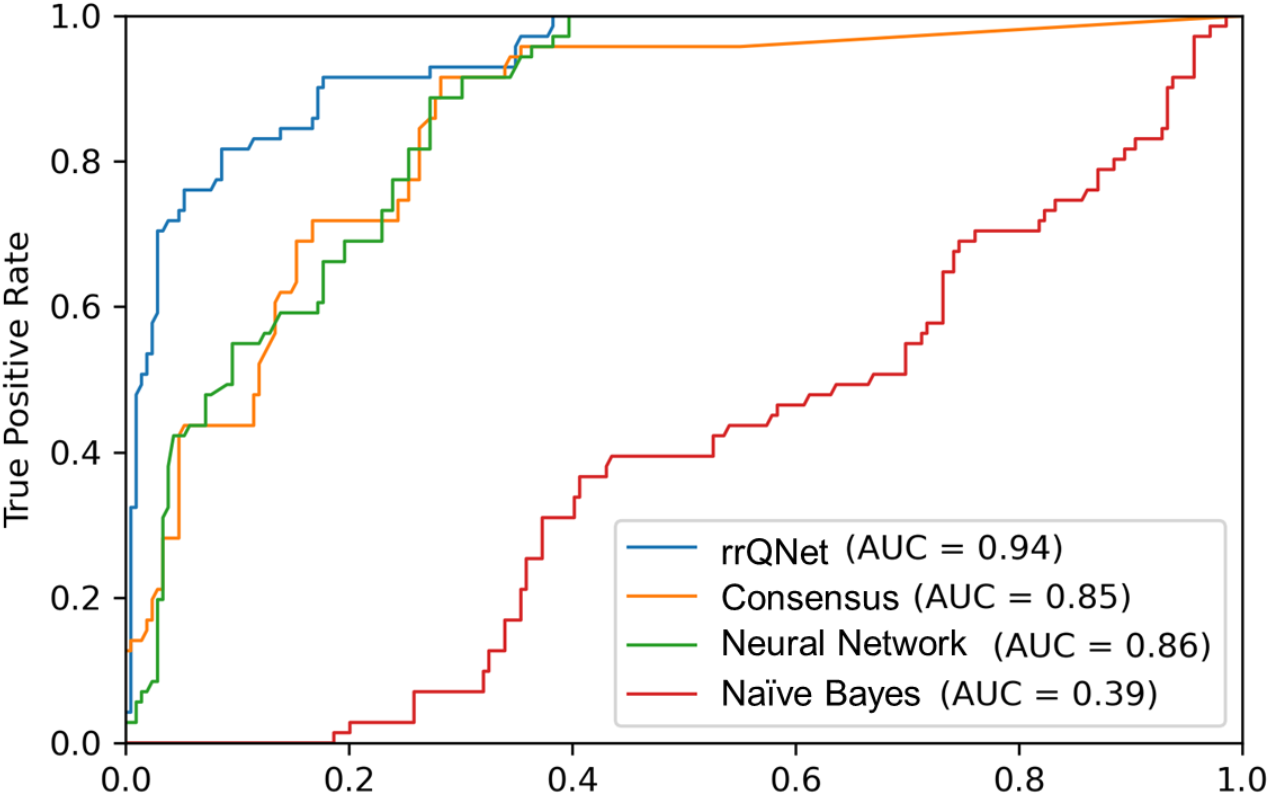
Receiver operating characteristic (ROC) analysis. The ROC curve and the corresponding AUC (Area Under Curve) value are used to assess rrQNet performance compared to our in-house baseline approaches implementing consensus, shallow neural network, and naïve Bayes in distinguishing between accurate and inaccurate contact prediction with a ground truth precision cutoff of 0.6 on 280 contact maps predicted by the seven locally installed contact prediction methods for 40 FM targets from CASP12 and CASP13.

### Confidence prediction of contacts at residue-pair resolution

While rrQNet ultimately provides an estimated quality score of a full-fledged input contact map, the penultimate stage of the rrQNet pipeline outputs a 2D matrix of predicted confidence score at the level of individual residue-pairs from the input contact map. That is, rrQNet intrinsically performs confidence prediction of individual contacts at the residue-pair resolution to classify a contact as true positive if the average predicted likelihood of the residue pair (i,j) and the residue pair (j,i) is greater than the threshold of 0.5. In **Figure 3** (A)-(F), we show the true positive contacts predicted by rrQNet (lower triangles, green dots) out of the top L predicted contacts (lower triangles, red dots) against the ground truth (upper triangle, blue dots) for six representative targets from the CASP12 and CASP13 benchmark sets. For targets T0864-D1, T0968s2-D1, and T0969-D1, the contact maps predicted by ResPRE, MetaPSICOV, and DeepConPred2, respectively, are shown as representative examples of accurate contact prediction. For each of the three contact predictions shown in Figure 3 (A)-(C), true positive contacts predicted by rrQNet (green dots) cover almost all the contact patterns present in the ground truth (blue dots). As such, the final estimated quality scores predicted by rrQNet for the three targets are 0.87, 0.74, and 0.7, respectively, which are close approximations of the corresponding ground truth precision scores of 0.9, 0.71, and 0.65, respectively. On the other hand, for target domains T0904-D1, T0950-D1, and T0980s1-D1 shown in Figure 3 (D)-(F), contact maps predicted by FreeContact and PSICOV are examples of noisy contact prediction, having ground truth precisions of 0.02, 0.01, and 0.09, respectively. Here, rrQNet correctly recognizes most of these contacts as false positives, leading to the final estimated quality scores of 0.03, 0.01, and 0, respectively. In summary, rrQNet is useful in predicting the confidence of contacts at the residue-pair resolution.

**Figure 3.**
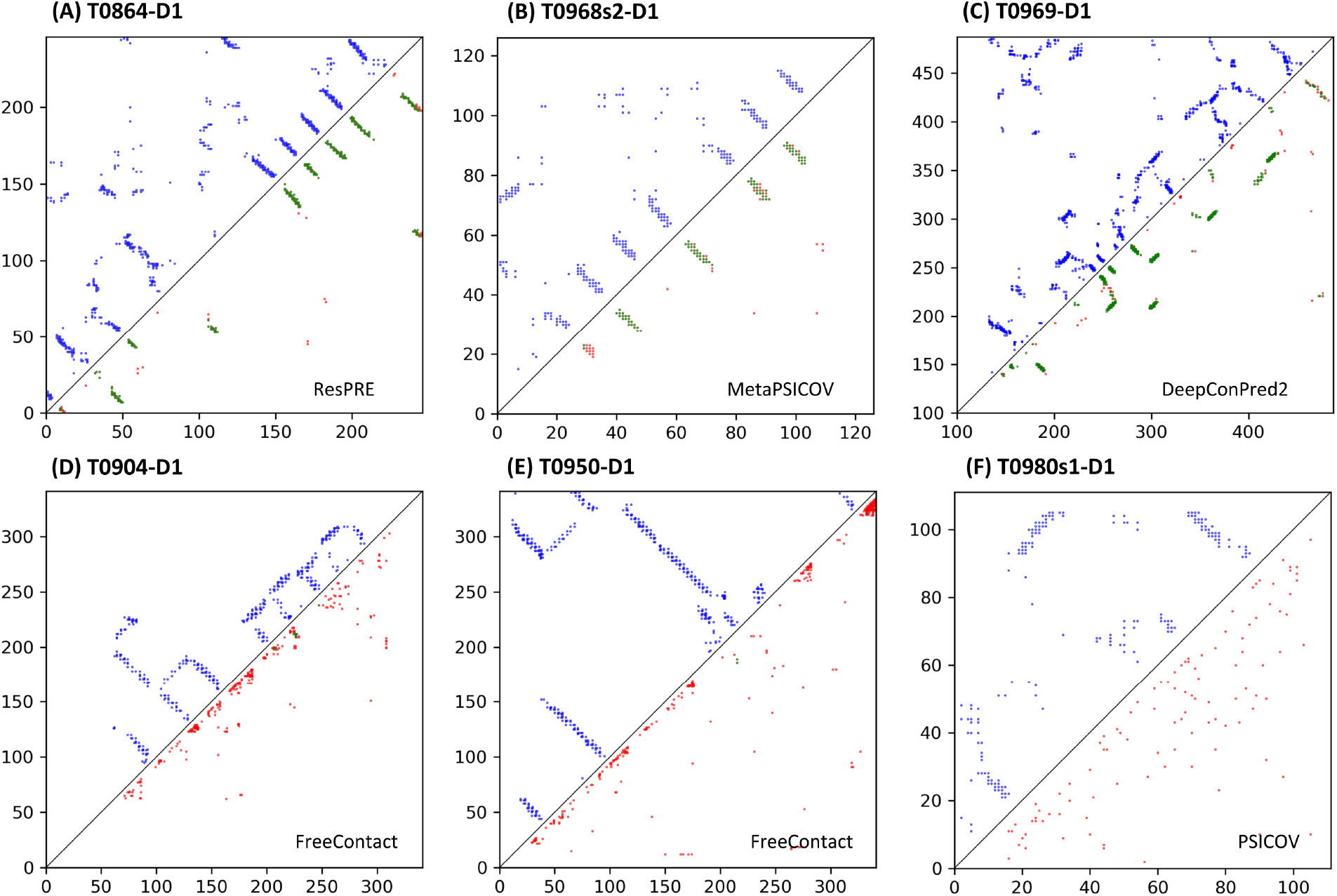
Confidence prediction of contacts. Six representative examples are shown. The upper triangle with blue dots represents the ground truth, the lower triangle with red dots represents the top L predicted contacts using various predictors, and the lower triangle with green dots represents the contacts classified by rrQNet as true positives.

To investigate whether the performance of rrQNet generalizes for the state-of-the-art contact and distance predictors such as AlphaFold2^5^, trRosetta^3^, and RaptorX^1^, we study rrQNet prediction at the residue-pair resolution resulting from AlphaFold2, trRosetta, and RaptorX. **Figure 4** (A)-(C) shows the performance of rrQNet at the residue-pair resolution on three representative targets from CASP12 and CASP13 benchmark sets. Similar to **Figure 3**, the upper triangle (blue dots) represents the ground truth, and the lower triangle represents the top L predictions (red dots) and true positive contacts predicted by rrQNet (green dots). For targets T0859-D1, T0866-D1, and T1022s1-D1, contact maps predicted by AlphaFold2, trRosetta, and RaptorX, respectively are shown as representative examples of high accuracy contact predictions. trRosetta and RaptorX contact predictions are obtained by summing up the likelihood values of the predicted distance bins up to 8Å, and AlphaFold2 contact prediction is obtained from the file ‘rank_1_model_1_ptm_seed_0.raw.txt’ by running AlphaFold2 in ColabFold’s^42^ notebook. For all three cases in Figure 4 (A)-(C), rrQNet (green dots) captures the majority of contact patterns that are present in the ground truth (blue dots). All three contact maps are accurately predicted having the ground truth precisions of 0.87, 0.98, and 0.92, respectively, and graph-based per-node strength correlations of 0.75, 0.76, and 0.81, respectively. The final estimated scores by rrQNet are also high as 0.92, 0.84, and 0.83, respectively. The results demonstrate the generalizability of rrQNet for confidence prediction of contacts at the residue-pair resolution for the state-of-the-art predictors, even when the train set of rrQNet does not include these methods.

**Figure 4.**
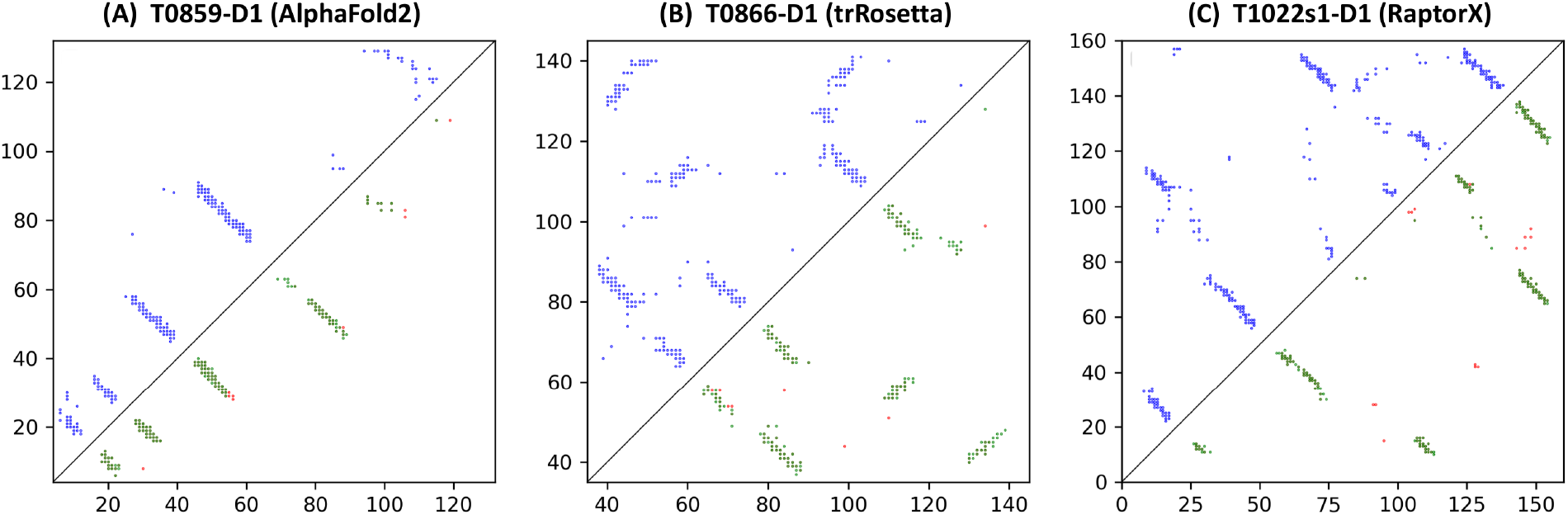
Confidence prediction of contacts predicted by AlphaFold2, trRosetta, and RaptorX. Three representative examples are shown. The upper triangle with blue dots represents the ground truth, the lower triangle with red dots represents the top L predicted contacts using various predictors, and the lower triangle with green dots represents the contacts classified by rrQNet as true positives.

### Pseudo-blind assessment on CASP14 contact prediction experiment

To investigate the generalizability of rrQNet in accurately estimating the qualities of contact maps beyond the seven locally installed contact prediction methods used for training and benchmarking, we perform a pseudo-blind assessment by estimating the qualities of the contact maps submitted by 30 server predictors participating in CASP14 contact prediction category. **Figure 5** shows the rrQNet estimated quality score versus the CASP reported ground truth precision, averaged over all targets submitted by various server predictors for long-range, medium- and long-range, and short-range contacts. rrQNet consistently achieves high positive correlations between the estimated contact map quality and ground truth precision. For instance, the Pearson correlation (r) for long-range only contacts is 0.83, combined medium- and long-range contacts is 0.88, and short-range only contacts is 0.92. Spearman correlation (ρ) (and Kendal’s Tau correlation (τ)) for long-, medium- and long-, and short-range contacts are 0.7 (and 0.53), 0.69 (and 0.53), and 0.7 (and 0.55), respectively. It is interesting to note that rrQNet continues to deliver good quality estimation performance for a wide variety of methods participating in CASP14, some of which are significantly improved versions of contact predictors than those used in rrQNet training. The results demonstrate that rrQNet can generalize well for estimating the qualities of contact maps predicted by a diverse array of methods. This suggests that the quality score of rrQNet can be potentially used as a loss function of an end-to-end structure prediction system as well as for iterative optimization of the contact maps by propagating the feedback through the end-to-end architecture. Furthermore, contact-driven homology detection and reconstruction approaches can benefit from such a generalizable quality estimate of a contact map predicted by external methods.

**Figure 5.**
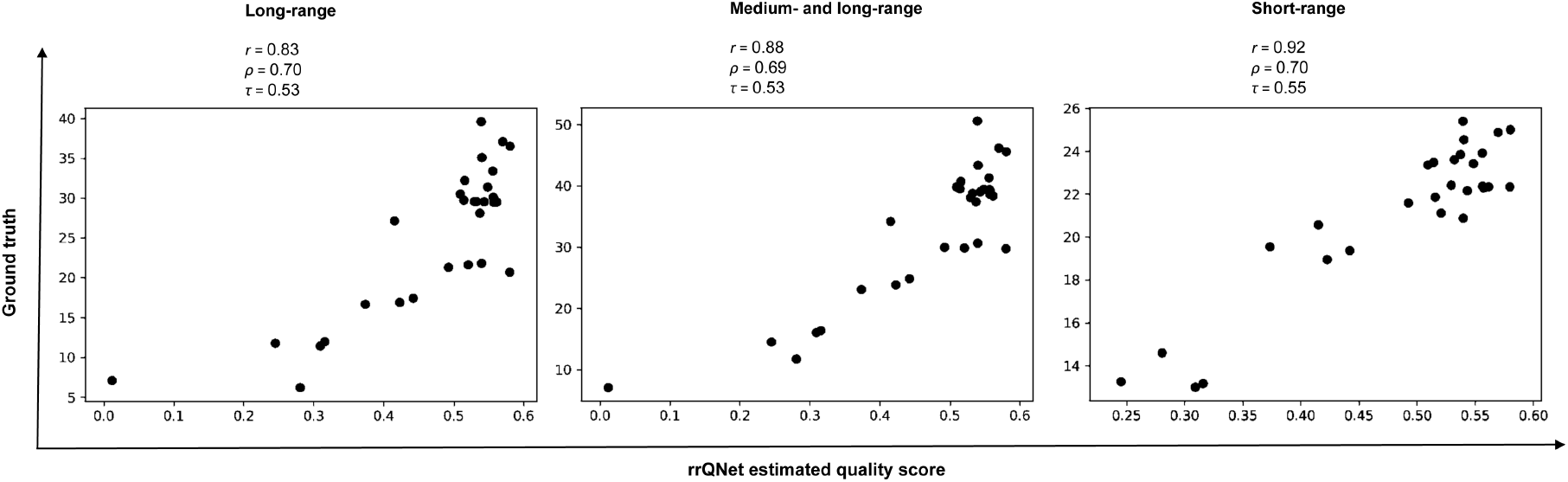
Estimated quality score versus ground truth precision for CASP14*. rrQNet estimated quality score versus the CASP reported ground truth precision, averaged over all targets submitted by various server predictors are shown for long-range, medium- and long-range, and short-range contacts. *BrainFold method is discarded for the calculation of short-range contacts due to missing prediction.

In **Figure 6** (A)-(F), we show the performance of rrQNet at the residue-pair resolution on contact maps predicted by different server predictors for three representative CASP14 targets. For targets T1047s1-D1, T1074-D1, and T1090-D1, contact maps predicted by MULTICOM-DIST, PrayogRealDistance, and tFold-Cat are examples of reasonably accurate predictions (Figure 6 (A)-(C)), whereas contact maps predicted by BrainFold and ICOS are noisy (Figure 6 (D)-(F)). For all the three high-quality predicted contacts, rrQNet (green dots) estimates the majority of contact patterns present in the ground truth (blue dots), leading to the final estimated quality scores of 0.9, 0.88, and 0.83, for MULTICOM-DIST, PrayogRealDistance, and tFold-Cat predictions, respectively, which are close approximation of their ground truth precisions of 0.88, 0.73, and 0.9, respectively. On the other hand, for noisy contact maps predicted by BrainFold and ICOS, rrQNet correctly recognize most of the false positives, resulting in the final estimated quality scores of 0, 0.02, and 0.37, respectively, which are close approximation of ground truth precision of 0.06, 0.13, and 0.39, respectively. It is interesting to note that rrQNet correctly recognizes the sparsely populated segment of accurate pattern in the contact map predicted by ICOS, despite the noise in the overall predicted contact map. The results demonstrate the robustness of rrQNet performance.

**Figure 6.**
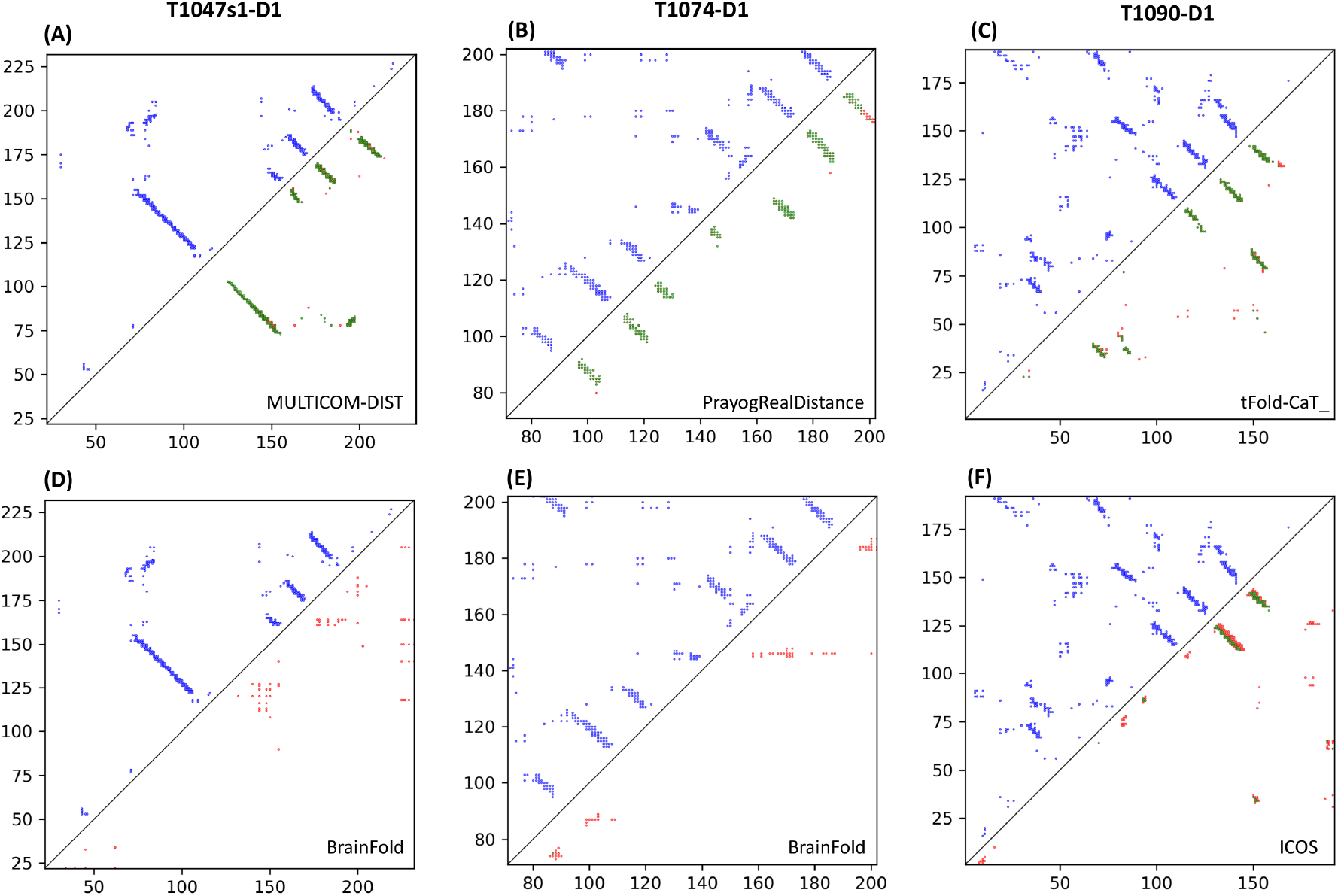
Confidence prediction of contacts predicted by different servers participating in CASP14. Six representative examples are shown for three targets. The upper triangle with blue dots represents the ground truth, the lower triangle with red dots represents the top L predicted contacts using various predictors, and the lower triangle with green dots represents the contacts classified by rrQNet as true positives.

### Ablation study

To examine the contribution of various components used in our deep learning-based contact map quality estimation method such as the deep neural network architecture adopted and the contact map prediction methods utilized for the training, we perform an ablation study using an independent validation set of 30 FM targets from CASP11. We follow the same multiple sequence alignment (MSA) generation protocol and the same training and inference procedure as described before to perform head-to-head comparison.

#### Contribution of deep neural network architecture

rrQNet consists of the evolutionary and reconciliation modules, each being a stack of residual neural network (ResNet) blocks. We study the effect of the network architecture on performance by gradually varying the number of ResNet blocks in the evolutionary and reconciliation modules while utilizing the same training dataset and evaluating quality estimation performance on the CASP11 validation set. As shown in **Figure 7**, the global Pearson correlation steadily increases with the increasing number of the ResNet blocks and attains a value of 0.8 for 55 ResNet blocks in the evolutionary module and 40 ResNet blocks in the reconciliation module (hereafter called the 55/40 architecture), before sharply dropping, possibly due to overfitting when deeper architecture beyond 55/40 is used. The per-target average Pearson correlation remains relatively steady while attaining the highest value of 0.91 for the 55/40 architecture. As such, this architecture is empirically chosen as the de facto model used in rrQNet. When we replace the ResNet blocks with standard convolutional blocks in the de facto model, the global Pearson correlation drops from 0.8 to 0.77, thus justifying the choice of ResNet architecture used in rrQNet.

**Figure 7.**
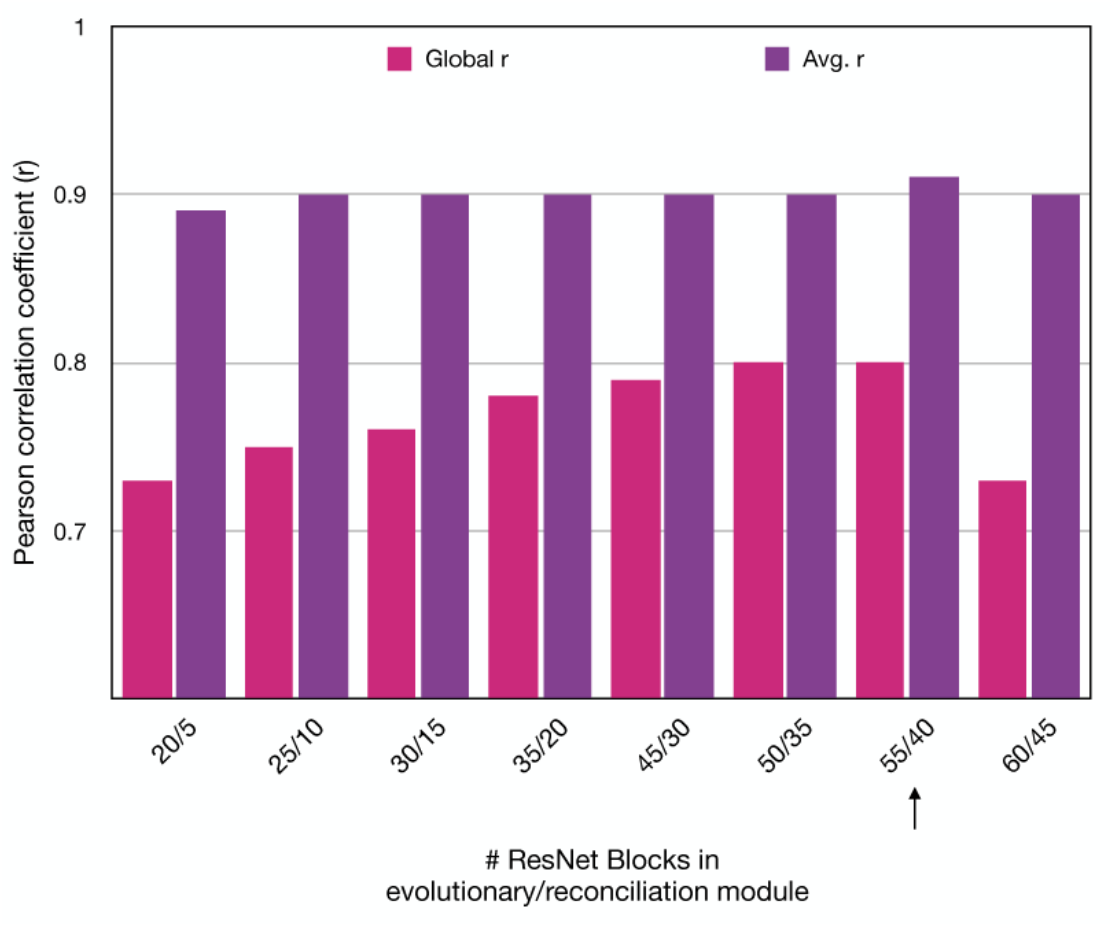
Effect of deep neural network architecture. Global and per-target average Pearson correlations are shown for a gradually increasing number of ResNet blocks in the evolutionary and reconciliation modules. The arrow indicates the *de facto* model chosen in rrQNet.

#### Contribution of contact prediction methods used for training

Our deep learning model is trained on seven contact predictors ranging from pure co-evolutionary methods to deep learning approaches. A natural question to ask is can we improve the performance even further by discarding the low-accuracy co-evolutionary contact prediction methods for training and instead exploiting cutting-edge approaches for predicting finer-grained binned distance distributions, which can lead to contact maps as a byproduct? To examine such question, we retrain rrQNet after discarding pure co-evolutionary methods PSICOV and FreeContact and evaluate quality estimation performance on the CASP11 validation set. The model retrained without PSICOV and FreeContact underperforms (global Pearson correlation 0.78) the original rrQNet (global Pearson correlation 0.8). We also independently retrain rrQNet after adding two recent distance map predictors trRosetta^3^ and RaptorX^1^ and converting the binned distance distributions into contact maps by summing up the likelihood values of the predicted distance bins up to 8Å. The model retrained with trRosetta and RaptorX still underperforms (global Pearson correlation 0.77) the original rrQNet (global Pearson correlation 0.8). The results suggest that pure co-evolutionary methods used in rrQNet are useful, possibly by calibrating the training data in terms of false positives, whereas finer-grained distance map predictors may not bring additional value for contact map quality estimation.

## DISCUSSION

We have introduced rrQNet, a deep learning method for contact map quality estimation that can accurately perform self-assessment of protein structures at the residue pair representation captured by a 2D contact map, without explicitly turning the contact map into 3D coordinates. By performing deep neural network-based reconciliation of the evolutionary context-integrated contact maps, our trained classifier distinguishes true positive from false positive contacts, ultimately approximating the precision of an input contact map. We have shown that its predictions achieve much better performance than our in-house baselines, generalize well for contact maps generated by a wide variety of contact predictors, and offer versatility for contact quality estimation at varying granularities from individual residue-pair to full-fledged contact map. The contact quality score predicted by our method may help improve end-to-end protein structure prediction.

We may further improve the sensitivity of our method by extending the deep learning model to identify false negative contacts, which can be probabilistically combined with the estimated true positives in order to estimate the recall of a predicted contact map. Estimated recall can be used together with the predicted precision for a more rigorous evaluation of the input contact map. This will likely improve the robustness of our method by approximating the F-score metric, which can be calculated as the harmonic mean of the estimated precision and recall values. We may also improve the quality estimation performance by combining both evolutionary and physical constraints. For example, we may enforce a set of realistic physical constraints on the contact map^43^, along with the recognition of protein-like contact patterns^44^. Such integration of both evolutionary and physical constraints using a deep learning model could result in improved quality estimation. Finally, instead of quality estimation of predicted contact maps, our deep learning model can be extended to estimate the quality of an inter-residue distance matrix, which encodes finer-grained information than contact maps and provides more physical constraints of a protein structure and thus, potentially benefits protein structure prediction more than the quality estimation of predicted contacts. In this regard, the quality estimation of the inter-residue distance matrix can include the evaluation of inter-domain accuracy in addition to domain-level correctness by incorporating AlphaFold’s Predicted Aligned Error (PAE) plot, in which each position (x, y) captures the expected distance error in residue x’s position when the prediction and true structure are aligned on residue y. This should be useful in estimating the correctness of relative domain orientations for large multi-domain protein structures.

## ACKNOWLEDGEMENT

This work was partially supported by the National Institute of General Medical Sciences [R35GM138146 to DB] and the National Science Foundation [IIS-2030722, DBI-1942692 to DB].

## Conflict of Interest

none declared.

## Notes

### Competing Interest Statement

The authors have declared no competing interest.

